# Bioelectricity generation by *Symbiodinium microadriaticum*: a symbiont-forming photosynthetic dinoflagellate alga from coral reefs

**DOI:** 10.1101/2025.05.16.654495

**Authors:** Loris Marcel, James T. Simon, Joshua M. Lawrence, Svetlana Menkin, Adrian C. Barbrook, R. Ellen R. Nisbet, Christopher J. Howe, Jenny Z. Zhang

## Abstract

Climate change is intensifying the phenomenon of coral bleaching, a highly ecologically damaging process that stems from the breakdown in the symbiotic relationship between photosynthetic dinoflagellate algae and Cnidaria (animals). Currently, little is understood about bleaching at the molecular level, since changes in metabolic and redox interactions between the dinoflagellate algae and Cnidaria are challenging to study. Here, we developed electrochemical approaches to measure extracellular electron transfer (EET, a type of bioelectricity) in *Symbiodinium microadriaticum*, a model species of symbiont-forming dinoflagellate algae, and show how this can be used to assess intracellular bioenergetic fluctuations within the alga. We show that EET is dependent on two major intracellular pathways – photosynthesis and respiration – and reveal that expelled electrons exit the cell via diffusible electron carriers. We confirm that EET activity can be affected by environmental stressors linked to coral bleaching, such as changes in temperature, pH and light intensity. Overall, this is a direct and non-invasive approach to provide quantitative measurements of *S. microadriaticum*’s bioenergetics and redox exchanges with the environment in stress conditions. This platform can aid our understanding of the coral-dinoflagellate symbiosis and the molecular mechanisms of bleaching.

## Introduction

Dinoflagellates are single-celled photosynthetic eukaryotes that can form symbiotic relationships with a range of animals, including corals (Fig. 1a-b); they provide corals with sugars produced through oxygenic photosynthesis and carbon fixation in exchange for nitrogen, phosphate and inorganic carbon^1,2^. In periods of environmental stress, such as increased water temperature, the coral-dinoflagellate symbiosis can break down^3^. Dinoflagellate algae are subsequently expelled from coral cells in a process called bleaching, which puts corals at risk of starvation. With climate change, coral bleaching events are becoming increasingly frequent, and corals may even become extinct by the end of the century, particularly in low-latitude regions^4^. Despite being essential members of reef ecosystems, symbiotic dinoflagellates are still poorly studied. This is mainly due to the difficulty of modifying dinoflagellates genetically^5^, although some success in nuclear and chloroplast transformation has been reported^6,7^. The difficulty of genetic modification may be due in part to the permanently condensed state of the nuclear genome^8^ and the fragmentary nature of the organelle genomes^9^.

**Fig. 1:**
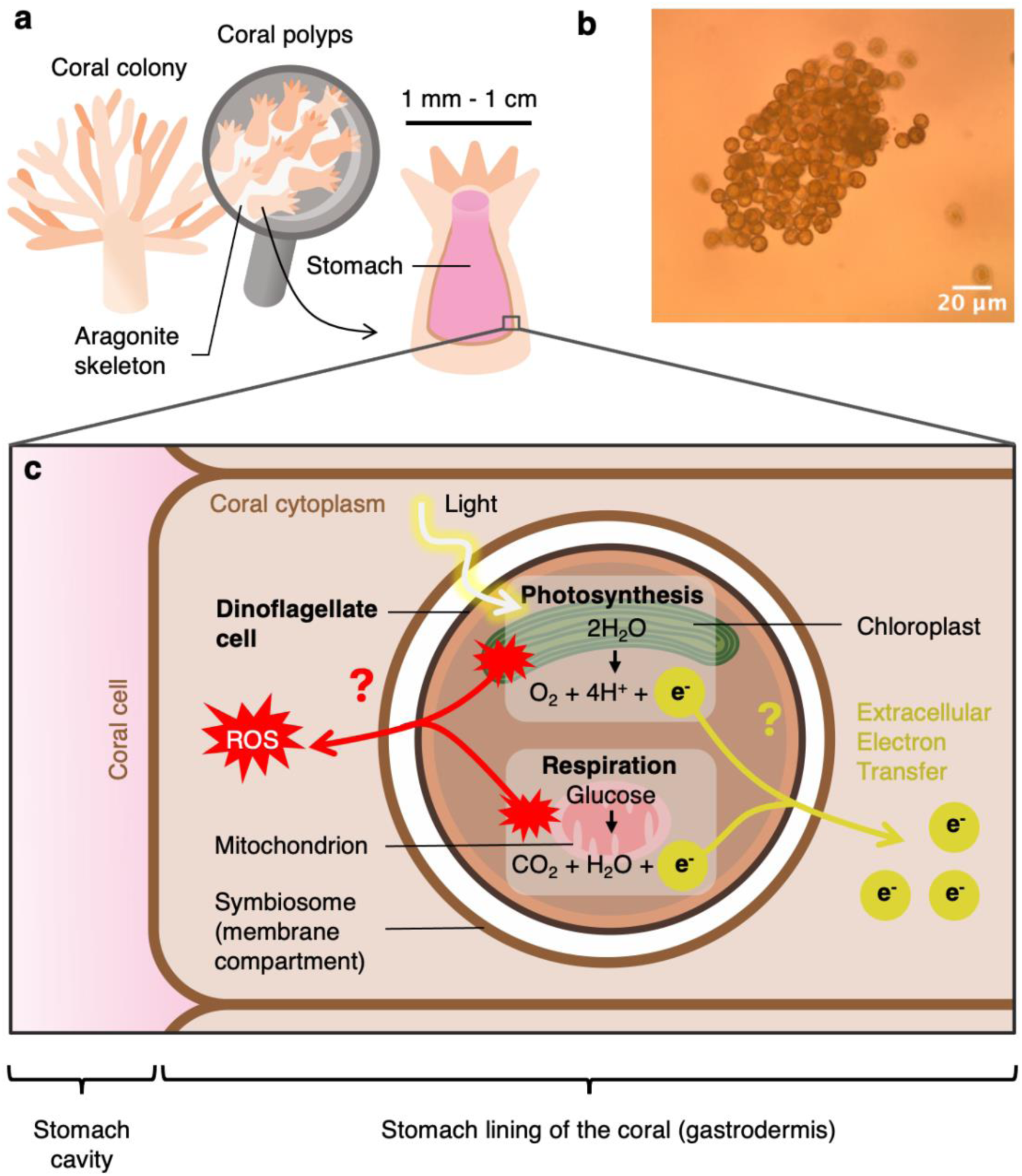
Interactions between corals and their dinoflagellate symbionts. **a** Anatomy of corals. A coral colony is composed of individual animals called polyps, which grow on a shared aragonite skeleton. Polyps contain a large stomach cavity, the lining of which is populated with symbiotic dinoflagellate algae together with other microorganisms (both bacteria and eukaryotes). **b** Bright-field microscopy image of an aggregate of the model dinoflagellate studied, *Symbiodinium microadriaticum*. **c** Simplified schematic of the redox exchanges that could occur between dinoflagellates and their coral hosts. Dinoflagellate algae are maintained in the cytoplasm of coral cells, so their photosynthetic and respiratory pathways are likely to interact with the coral intracellular environment. Abbreviations: ROS, reactive oxygen species; e^-^, electrons.

Many photosynthetic microorganisms can export electrons out of their cell in a bioelectricity-generating process termed extracellular electron transfer (EET), for which several possible mechanisms have been proposed^10–12^. Heterotrophic prokaryotes may use EET to reduce extracellular terminal acceptors to drive metabolism^11,13^, or to facilitate syntrophy via interspecies electron transfer^14^. The functions of EET in photosynthetic organisms have yet to be conclusively demonstrated, although EET is linked to intracellular bioenergetic processes^12^. For example, in cyanobacteria, EET rate has been shown to be influenced by both photosynthetic and respiratory electron transport activities^12,15–17^. As eukaryotic algae with a chloroplast, dinoflagellates have a similar photosynthetic electron transport chain (PETC) to cyanobacteria. They harvest light energy to power a water-oxidation reaction in which electrons are extracted from water (Fig. 1c). These electrons flow through a series of proteins in the PETC, ultimately producing the energy-carrying molecule adenosine triphosphate (ATP) and the redox carrier nicotinamide adenine dinucleotide phosphate (NADPH) which can be used to fuel cellular metabolic processes. Some of the electrons generated during photosynthesis can leave the cell via the EET pathway, generating bioelectricity.

Photosynthesis may also generate reactive oxygen species (ROS) which play an important role in cell signalling and can result in oxidative damage to a variety of biological molecules (Fig. 1c).

We hypothesized that dinoflagellates can carry out EET. If they do, it would potentially facilitate metabolic interactions with coral which could assist or regulate the symbiosis (Fig. 1c). Additionally, as bleaching events are often correlated with algal redox stress^18–23^, the ability to measure the extracellular and intracellular redox balance via EET in dinoflagellates would be a particularly useful tool for assessing coral health. However, it is currently unknown whether – or how – dinoflagellates perform EET (Fig. 1c).

Here, we develop an electrochemical platform to measure EET in the important symbiont-forming dinoflagellate species *Symbiodinium microadriaticum*. We report for the first time that *S. microadriaticum* performs EET, we demonstrate the dependence of EET on intracellular bioenergetics, and we show that EET is responsive to stress conditions known to induce coral bleaching. We discuss the implications of dinoflagellate EET in the context of the coral-dinoflagellate symbiosis and for the development of coral diagnostics.

## Results

### Developing an electrochemical platform for analysing EET in *S. microadriaticum*

EET from photosynthetic microorganisms can be measured analytically in three-electrode electrochemical systems. These systems contain a working electrode, where the redox reaction of interest can be studied and compared against a well-characterised reference electrode. Typically, cells can be adhered to the working electrode to enable sensitive measurements of the biological current resulting from metabolic activity. In the case of photosynthetic cells, illumination causes an increase in this current, termed a photocurrent, which is caused by an enhancement of EET due to increased photosynthetic electron transfer activity^12,15^. Measuring photocurrents provides a readout of photosynthetic activity as well as other pathways that may contribute to EET.

We designed a three-electrode system to measure EET in *Symbiodinium microadriaticum*, a model species of symbiotic dinoflagellates^24^ (Fig. 2a). A horizontal working electrode was utilised (Fig. 2a-b, Supplementary Fig. 1a-c) because *S. microadriaticum* readily settles by forming aggregates (Fig. 1b). Mesoporous indium-tin oxide (mesoITO) was used as a working electrode due to its high surface area, biocompatibility and translucency^25^. *S. microadriaticum* cells absorb strongly in red and blue light (Fig. 2c) due to the presence of chlorophylls and carotenoids involved in photosynthesis^26^. A 680 nm light source was chosen to trigger photocurrents because ITO can produce abiotic currents in blue light^27^.

**Fig. 2:**
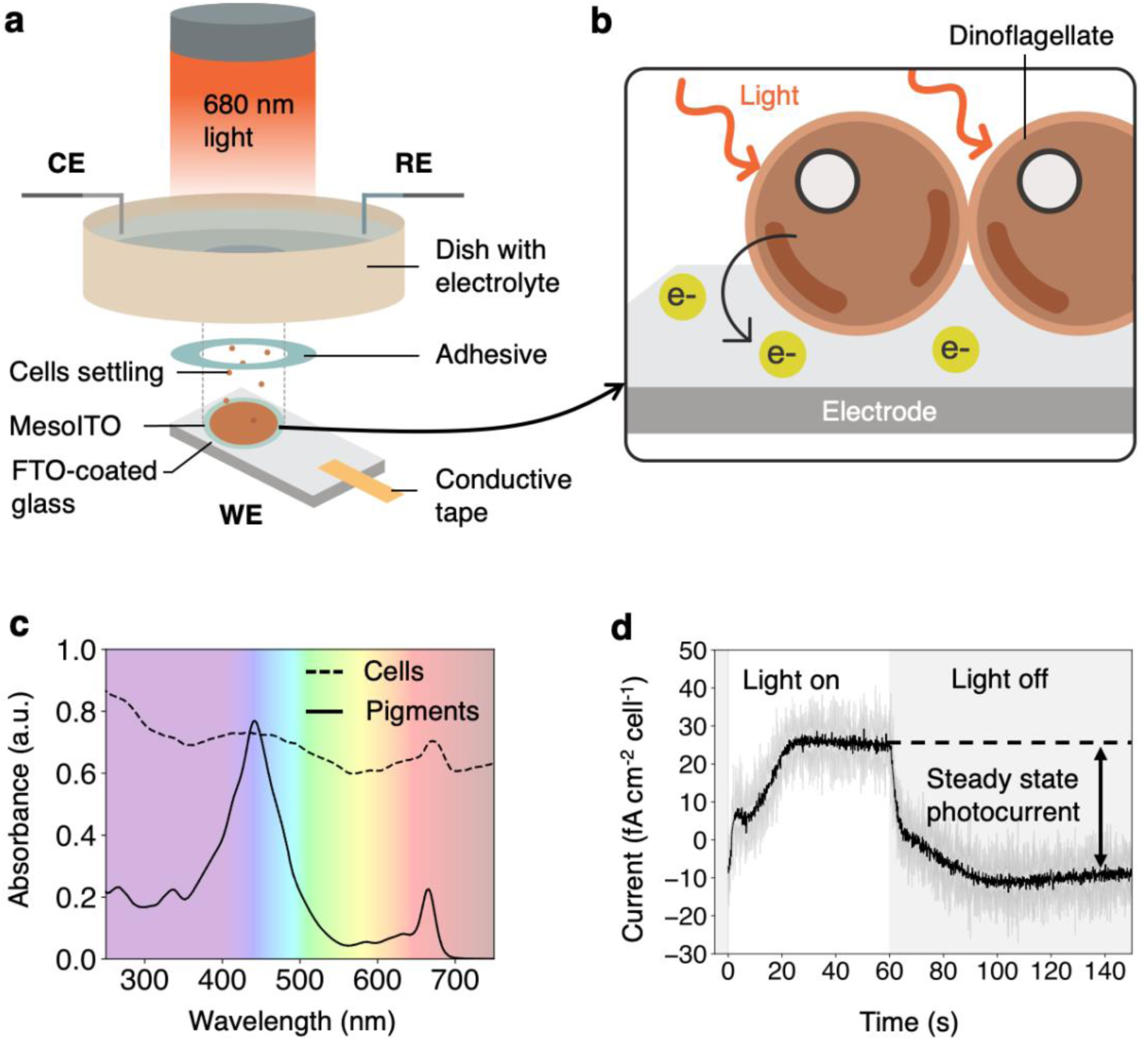
An electrochemical platform to study *S. microadriaticum*. **a** Schematic of the electrochemical platform used in this study. *S. microadriaticum* cells were settled onto a horizontal mesoITO electrode, which acted as the working electrode (WE) in a three-electrode system. An Ag/AgCl electrode and a platinum mesh were used as the reference and counter electrodes (CE and RE) respectively. **b** Schematic of EET in dinoflagellates in which electrons leave the cell and reduce the electrode. **c** Absorbance spectrum of *S. microadriaticum* cells and their pigments extracted with methanol and ammonium acetate. **d** Photocurrent profile (current shape) obtained from *S. microadriaticum*. The steady state photocurrent is the magnitude of the current, defined by difference between the steady state currents in the light and in the dark. The photocurrent trace is the average of three biological replicates containing five technical replicates each, the shaded trace corresponds to standard deviation. Measurements were obtained at 0.3 V vs SHE; with 680 nm light at 50 μmol photons m^-2^ s^-1^; and 3 x 10^6^ cells.

Using the three-electrode system, a photocurrent of ∼25 fA cm^-2^ cell^-1^ was obtained (Fig. 2d). This was unexpectedly large, since *S. microadriaticum* has a thick cellulosic shell and a plasma membrane decorated by various glycans^28,29^ which could limit the diffusion of electron carriers out of the cell. The steady state photocurrent obtained with *S. microadriaticum* was 2.5-fold higher than reported for diatoms (which are also eukaryotic algae) without electron mediators^30^ – one difference was that 680 nm light was used here instead of white light.

The bio-electrochemical system was further optimised by testing the effect of cell loading on photocurrent production (Supplementary Fig. 1d). Maximal steady state photocurrents were obtained between 3 and 6 million *S. microadriaticum* cells per 1 cm-diameter electrode (0.785 cm^2^). The steady state photocurrent was reached faster with 3 million cells per electrode (Supplementary Fig. 1e), probably because larger cell numbers caused more shading of cells closest to the electrode. Hence this value was chosen for further experiments. Additionally, we found that *S. microadriaticum* cells were fully settled onto the electrode after 15 min (Supplementary Fig. 1f). *S. microadriaticum* in mid-exponential phase produced higher and more consistent currents than cells in lag phase or early stationary phase (Supplementary Fig. 1g).

Using a range of applied potentials, we showed that the earliest potential where steady state photocurrents became positive was 0.2 V vs the standard hydrogen electrode (SHE) (Supplementary Fig. 2a), although photocurrent profile features could be observed at lower potentials (Supplementary Fig. 2b). Below 0.2 V vs SHE, the reduction of photosynthetically-produced oxygen at the electrode produced large negative currents which may have hidden positive photocurrents. Oxygen could not be removed, even with initial purging with N_2_ and enzymatic oxygen removal systems^31^.

### EET is dependent on intracellular bioenergetics of *S. microadriaticum*

In dinoflagellate algae, there are two main energy-producing pathways – photosynthesis and respiration – which are localised within the chloroplasts and mitochondria respectively (Fig. 1c). To test whether EET would be a good reporter of intracellular bioenergetics in dinoflagellates, photocurrents were measured in the presence of inhibitors and substrates of these pathways.

First, photocurrents were measured in increasing conditions of 3-(3,4-dichlorophenyl)-1,1-dimethylurea (DCMU) – an inhibitor of photosystem II (PSII) which blocks plastoquinone reduction at the Q_B_ pocket of PSII, one of the first steps of the PETC (Fig. 3a). With increasing concentrations of DCMU, the steady state photocurrent decreased drastically, until it was fully abolished between 10 and 100 μM DCMU (Fig. 3b).

**Fig. 3:**
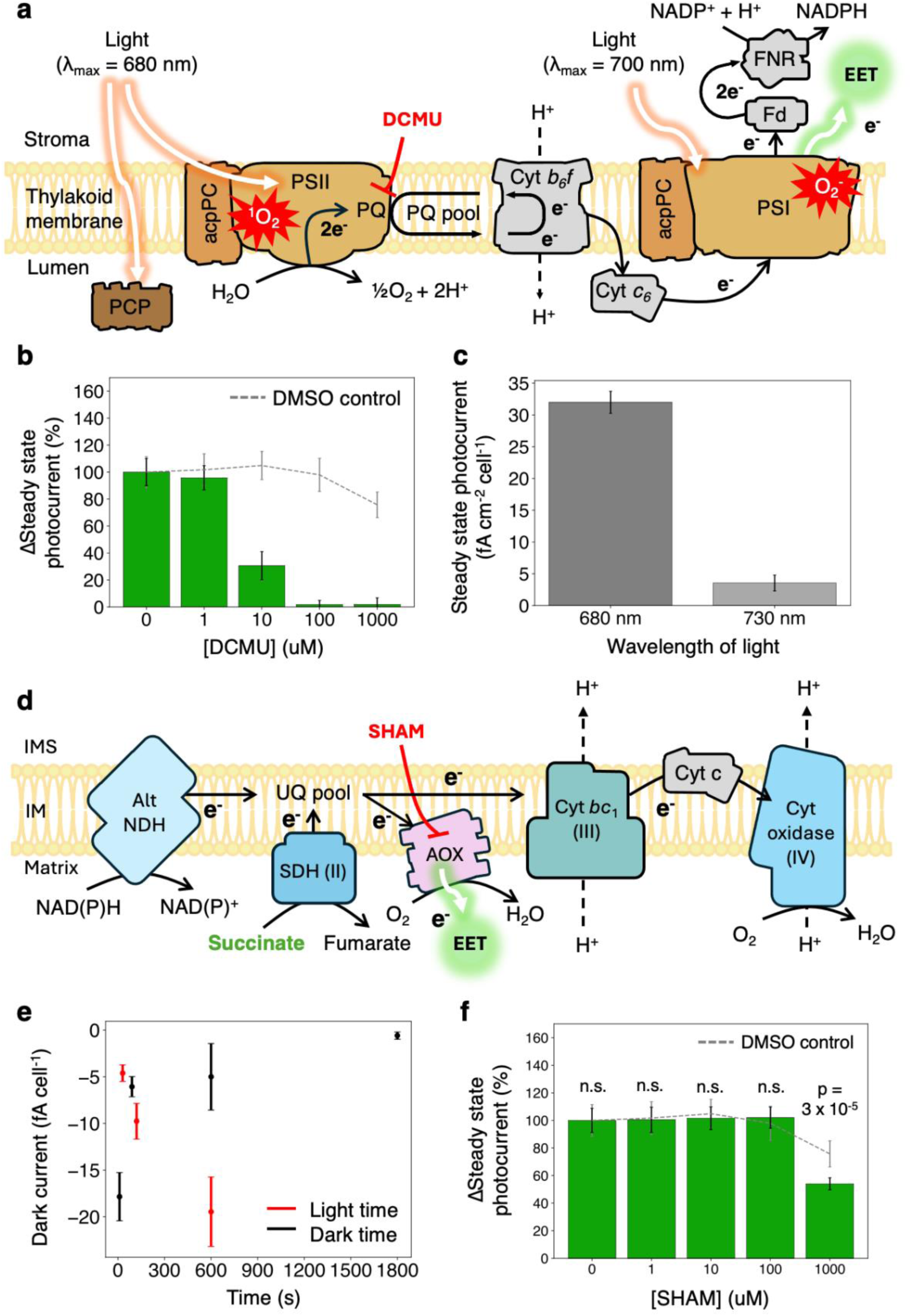
Extracellular electron export is dependent on intracellular bioenergetics. **a** The photosynthetic apparatus in *Symbiodinium* dinoflagellates. Abbreviations: PCP, Peridinin-chlorophyll *a* protein; acpPC, chlorophyll *a*-chlorophyll *c*_2_ peridinin protein complex; PQ, plastoquinone; Fd, ferredoxin; FNR Ferredoxin-NADP+ reductase; e^-^, electron; NADP+, nicotinamide adenine dinucleotide phosphate, ROS, reactive oxygen species. **b** Effect of DCMU on the steady state photocurrent. **c** Effect of 680 nm and 730 nm light on the steady state photocurrent. **d** Schematic of the respiratory electron transport chain in *Symbiodinium* dinoflagellates. Abbreviations: Alt NDH, alternative NADH dehydrogenase; SDH, succinate dehydrogenase; AOX, alternative oxidase; SHAM, salicylhydroxamic acid; Cyt, cytochrome; IMS, intermembrane space; IM, inner membrane. **e** Steady state dark currents from light/dark cycles where either the light period was varied and the dark period was kept at 90 s, or where the dark period was varied and the light period was kept at 60 s. **f** Effect of SHAM on steady state photocurrent. Data shown are averages of three biological replicates containing five technical replicates each, error bars represent the standard error of the mean. P-values were calculated with a paired samples t-test. Unless specified, measurements were obtained at 0.3 V vs SHE; with 680 nm light at 50 μmol photons m^-2^ s^-1^; and 3 x 10^6^ cells.

Previously, it was shown that PSII activity could be inhibited in *Symbiodinium* sp. cells at DCMU concentrations ranging from 20-80 μM, both in free-living cells^32^ and in anemone hosts^33^. This suggests that the steady state photocurrent is dependent on PSII activity. Additionally, steady state photocurrents were found to be 10-fold higher in 680 nm light – which activates both PSII and PSI – than in 730 nm light, which mainly activates PSI^34^ (Fig. 3a, c). This shows that exported electrons may leave the PETC downstream of PSI, but originate from water oxidation at PSII (Fig. 3a).

Next, dark currents were measured in different light and dark incubation times. In the dark, photosynthetic electron transfer is inactive, so dinoflagellate algae obtain most of their reducing power from respiration which oxidises stored carbon-containing compounds to generate electrons. An increase in dark current would suggest that more electrons from respiration are being fed into the EET pathway. Dark current was higher in longer dark incubation times (with light incubation times kept constant), when respiration is expected to be upregulated, showing that respiration can contribute to EET activity. Conversely, in longer light incubation times (with dark incubation times kept constant), dark currents were more negative (Fig. 3e). This is consistent with photosynthesis being the predominant pathway in light, and a decrease in respiratory activity. In all cases, the dark current was negative, indicating the reduction of redox species at the electrode interface. Some of these redox species were present in the culture medium, as shown by the negative current obtained with medium only (Supplementary Fig. 3a), but some could also have been produced by the cells.

Photocurrents were measured in the presence of salicylhydroxamic acid (SHAM, 0 – 1000 μM), an inhibitor of the alternative oxidase (AOX, Fig. 3d) which accounts for up to 26% of respiratory activity in dinoflagellate algae^35^. SHAM only caused a decrease in EET from 1 mM (Fig. 3f), consistent with previous studies in which mM-range concentrations were needed to inhibit AOX-mediated respiration^35^. Additionally, addition of succinate, the substrate of complex II in the RETC (Fig. 3d), induced a decrease in EET activity (Supplementary Fig. 3b). Since succinate is also a substrate in the tricarboxylic acid cycle, the decrease in steady state photocurrent could be explained by the activation of anabolic pathways at higher concentrations of succinate^36^. This would draw reducing equivalents such as NAD(P)H into biosynthesis rather than respiration. Photocurrents were also measured in the presence of rotenone, an inhibitor of the canonical mitochondrial complex I. Complex I catalyses the first step of the RETC and oxidises NADH. Although it is not clear if the canonical complex I is present in dinoflagellate algae (and a type II NADPH-dehydrogenase is believed to be present)^37,38^, rotenone slightly reduced the steady state photocurrent (Supplementary Fig. 3c), suggesting some effect on a canonical complex I or a possible inhibitory effect on other respiratory or photosynthetic proteins.

Together, these experiments suggest that EET and respiration are linked, with photosynthesis generating substrate molecules that are oxidised by the mitochondria and with some of the electrons subsequently fed into the EET pathway. This is consistent with previous reports of energetic coupling between photosynthesis and respiration found in *Symbiodinium* species^39^.

### EET in *S. microadriaticum* is mediated by an endogenous diffusible redox molecule

In all electroactive organisms studied so far, electron export to the electrode was found to be either direct (via the direct contact of a protein or nanowire), or mediated by a diffusible species^11^. To test which mode of electron transfer was involved for *S. microadriaticum*, an ultra-microelectrode (UME) operated by a scanning electrochemical microscope (SECM) platform was used. SECM platforms have been used previously to study redox activity from biological cells, including photosynthetic microorganisms^15^ and plant thylakoids^40^. Here, we made use of a three-electrode system in which the working electrode was a 10 μm-diameter platinum tip (Fig. 4a). The main advantage of SECM in the study of EET is that the working electrode can be precisely positioned in space using a piezo stage (Fig. 4a). This allows comparison of the redox activity both near and far from the dinoflagellates (Fig. 4a), and can therefore be used to investigate indirect EET mechanisms unlike conventional three-electrode systems. Additionally, UMEs are very sensitive and can detect redox molecules in femto-molar concentrations^41^.

**Fig. 4:**
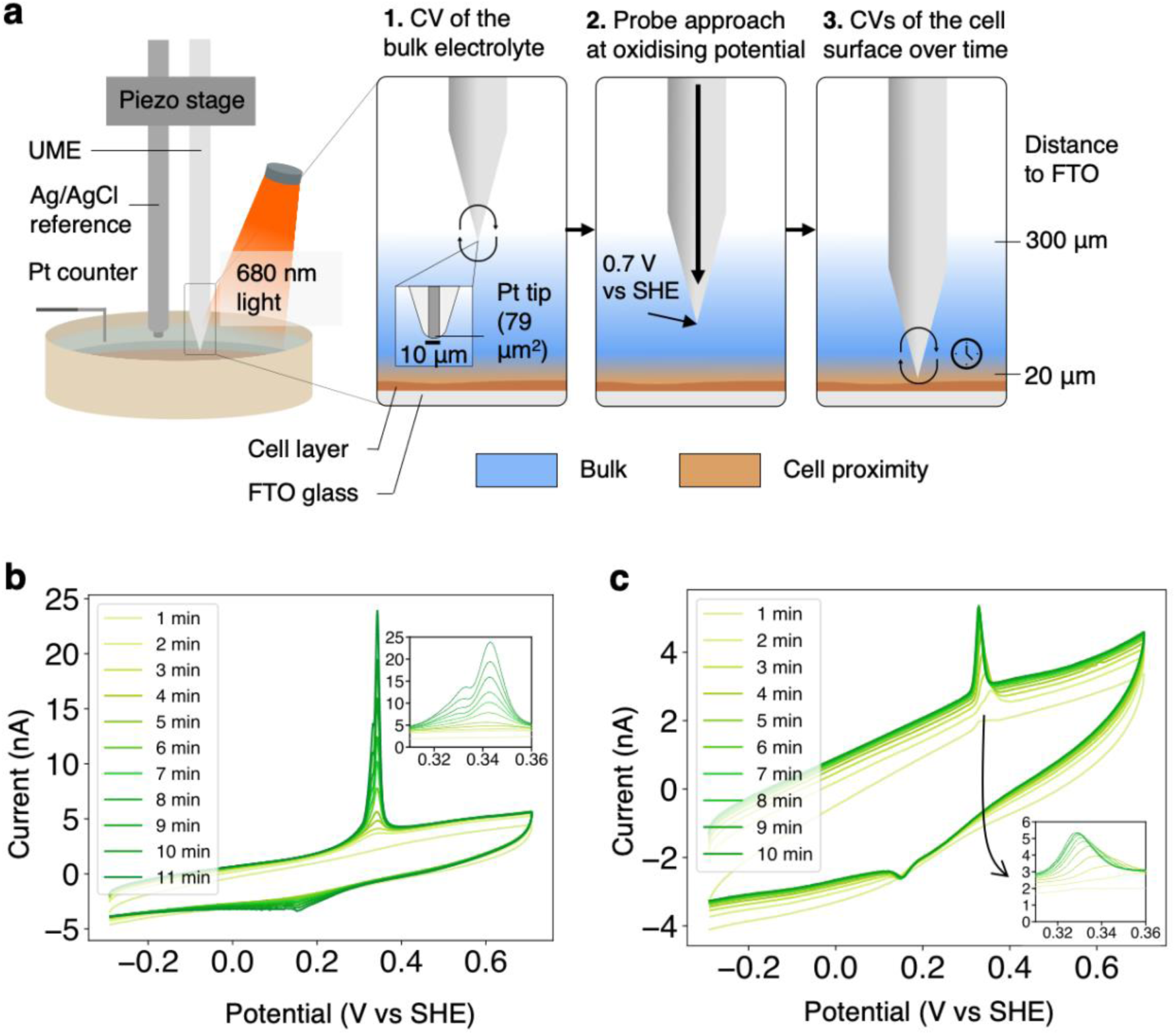
*S. microadriaticum* produces a diffusible electron carrier. **a** Schematic of the scanning electrochemical microscope (SECM) setup. The SECM is a three-electrode system where the working electrode is an ultra-microelectrode (UME) of diameter 10 μm. After two hours of incubation with or without 680 nm light at 100 μmol photons m^-2^ s^-1^, *S. microadriaticum* cells were subjected to three measurements: (1) a cyclic voltammogram (CV) of the bulk, (2) a probe approach curve at 0.7 V vs SHE and (3) a CV of the cell surface. **b** Representative CVs of the cell surface after 2h of incubation in light and a probe approach. **c** Representative CVs of the cell surface after 2h of incubation in the dark and a probe approach. CV experiments were repeated 3 times. All CVs were performed at 50 mV s^-1^ with a sample interval of 1 mV.

The presence of redox species on the UME was assayed with cyclic voltammetry (CV). In CV, the potential at the working electrode is cycled, and redox species present on the electrode surface may be oxidised or reduced, resulting in a current. In culture medium without cells, no redox activity was detected (Supplementary Fig. 4a). Likewise, when *S. microadriaticum* cells were added to fresh medium, there was no redox response in the bulk electrolyte (Supplementary Fig. 4b) or at the cell surface (Supplementary Fig. 4c). After 2 h of incubation in either 100 μmol photons m^-2^ s^-1^ or darkness, an oxidation peak at 0.33 V vs SHE was detected in the bulk, suggesting that a redox species was oxidised by the probe 300 μm away from the *S. microadriaticum* cells (Supplementary Fig. 4d).

The presence of the redox species in the dark is consistent with cells producing currents from respiratory activity, but the higher concentration of redox species after illumination shows that currents are mainly generated from photosynthetic activity. The spatial distribution of the detected redox species was then probed by performing CVs at the cell surface at regular time intervals. The concentration of the redox species increased as the measurements were performed, both in the light and the dark (Fig. 4b-c). Growth of the oxidation peak was greater in the presence of light, suggesting synthesis or secretion of this electron carrier is dependent on photosynthetic metabolism. This is supported by the complete inhibition of the oxidation peak in the presence of 100 μM DCMU (Supplementary Fig. 4e). The distance of this electron carrier from the cell surface, the dependence of its concentration on photosynthetic activity, as well as the fact its oxidation potential (0.33 V vs SHE) is close to the potentials at which maximal photocurrents were observed in experiments with mesoITO (> 0.2 V vs SHE, Supplementary Fig. 2a), all suggest that this electron carrier is responsible for EET in *S. microadriaticum*. In the light, the oxidation peak was split into two subpeaks (Fig. 4b). This could suggest that more than one electron carriers are produced, that the electron carrier is being converted into a different species, or that the electron carrier undergoes a sequential 2-electron oxidation process.

A reduction peak was also observed at 0.17 V vs SHE, although this was notably smaller in area than the oxidation peak, and the separation between the reduction and oxidation peaks was larger than expected for a reversible redox process (Fig. 4b-c). This suggests the two peaks do not correspond to the same analyte, with each analyte undergoing irreversible oxidation or reduction at the surface of the UME. Therefore, the analyte giving rise to the reduction peak is unlikely to influence EET, whilst the rate of EET is expected to be limited by the rate of synthesis of the analyte which gives rise to the oxidation peak. A control with no cells (without N_2_ purging) did not present any peaks after light incubation (Supplementary Fig. 4f), demonstrating the redox species giving rise to both the oxidation and reduction peaks were of biological origin and not due to degradation of f/2 medium components.

Overall, the results suggest that *S. microadriaticum* performs mediated EET. The electron carrier is diffusible, irreversibly oxidised by the electrode, and is produced in both light and dark albeit at different rates. While the concentration of the electron carrier cannot currently be calculated accurately, it is likely to be very low. The SECM probe typically yields Nernstian CV shapes because of its very small geometric area^42,43^, as shown by the measurements of the synthetic electron carrier FcMeOH at 1mM concentration (Supplementary Fig. 4g). However, the electron carrier detected here displays diffusion-limitation behaviour (Fig. 4b-c), suggesting that it could be present in low concentrations, or that its diffusion coefficient is low. Assuming that the electron carrier’s diffusion coefficient is similar to that of an amino acid^44^, we estimated the concentration of the electron carrier to be in the pico-molar range using the Randles-Sevcik equation (Supplementary Fig. 5).

### Tracking changes in EET in stress conditions known to induce coral bleaching

To explore the use of the electrochemical platform for diagnostic applications, we measured photocurrents in different stress conditions linked to bleaching. We hypothesised that the steady state photocurrent would vary in response to environmental changes.

First, we tested the effect of light stress. *S. microadriaticum* cells were kept in the dark for 1 h and were subsequently exposed to a series of light intensities to cause light stress. The light intensities ranged from 50 μmol photons m^-2^ s^-1^ (which represents typical growth conditions) to 1000 μmol photons m^-2^ s^-1^, which is higher than light intensities previously used to induce bleaching in corals^45^ (although 680 nm light was used here instead of white light). *S. microadriaticum* cells were subjected to the light stress regimes either in increasing or decreasing order of intensity. Oxygen evolution was also measured using an oxygen sensor. With light stress (decreasing), a markedly higher steady state photocurrent was measured at all light intensities (Fig. 5a). This confirms the coupling between EET and photosynthesis, and suggests that EET is responsive to environmental stress. A similar pattern was observed for oxygen production, although at 50 and 100 μmol photons m^-2^ s^-1^, EET increased proportionately more than oxygen evolution in the regime of decreasing light compared to the regime of increasing light (Fig. 5a-b). Since oxygen evolution can be used as a proxy for PSII activity, this result suggests that EET and photosynthesis are not fully coupled. Whilst this could provide evidence that EET sources electrons from downstream of PSII in the PETC (Fig. 3a) or from respiratory electron transport (Fig. 3d), these differences could also be explained by altered synthesis or secretion of electron carriers.

**Fig. 5:**
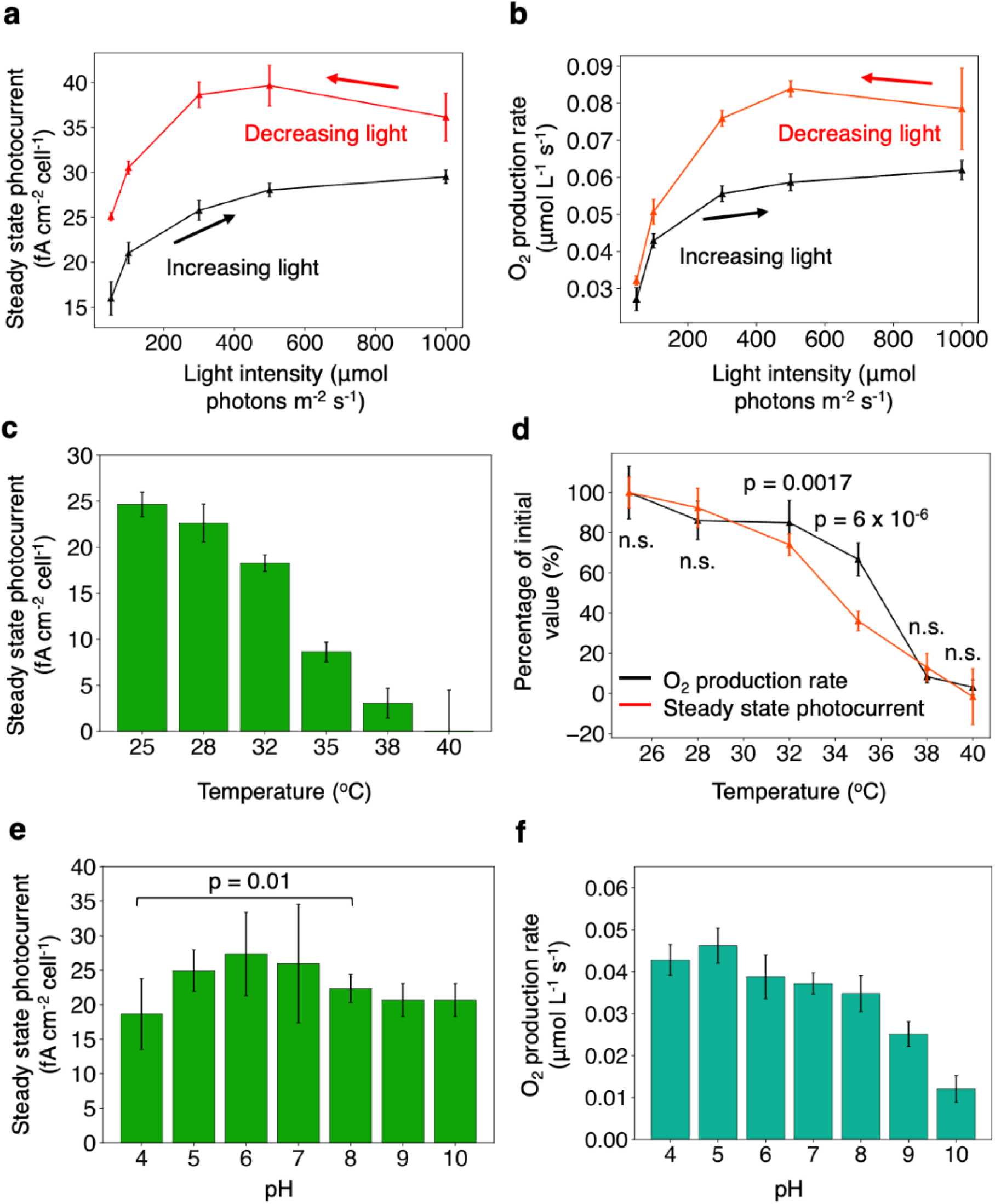
Extracellular electron export is responsive to changes in the environment. **a** Effect of light intensities ranging from 50 to 1000 μmol photons m^-2^ s^-1^ in increasing or decreasing order after incubation in the dark for 1 h. **b** Effect of the same light regimes as in *a* on oxygen production. **c** Effect of acute temperature stress on steady state photocurrent. Temperature was kept constant for 15 min during each photocurrent measurements, then ramped up at 1.5°C min^-1^ to the next temperature. **d** Comparison of the relative decrease in EET *vs* oxygen production in the same conditions. **e** Effect of acute pH stress on steady state photocurrent. *S. microadriaticum* cells were centrifuged and resuspended directly from pH 7.8 to the measured pH. **f** Effect of acute pH stress on oxygen production rates. Data shown are averages of three biological replicates containing five technical replicates each, error bars represent standard error of the mean. Statistics were performed using unpaired samples t-tests. Unless specified, measurements were obtained at 0.3 V vs SHE; with 680 nm light at 50 μmol photons m^-2^ s^-1^; and 3 x 10^6^ cells.

Additionally, we measured photocurrents at different temperatures since heightened water temperature is one of the main triggers of coral bleaching^3,46^. Steady state photocurrent density was strongly affected by heat stress, decreasing on average by 4%/°C between 25°C and 32°C (Fig. 5c). Growth rates of cell cultures also decreased with increasing temperatures (Supplementary Fig. 6a), suggesting a link between EET and cell health. However, the pH of the culture medium did decrease slightly at higher temperatures, which may have affected the cells (Supplementary Fig. 6b). While oxygen production and steady state photocurrent density both decreased with temperature, steady state photocurrent density showed a more direct correlation with temperature and was relatively more affected than oxygen evolution at 32°C and 35°C (Fig. 5d), which is further evidence of uncoupling between PSII activity and EET.

In the literature, pH stress was reported to impact coral calcification rates^47^ but not to cause bleaching directly, save for one report^48^ – in that study, acidification was found to cause bleaching after 8 weeks of treatment, but alkalinisation was not tested. We measured photocurrents at different artificially adjusted pHs, hypothesising that there would be no strong effect on steady state photocurrent. Whilst there was a significant difference in steady state photocurrent pH between pH 4 and pH 8 (the latter corresponding to the average seawater value), *S. microadriaticum* still produced detectable photocurrents at pH 4 (Fig. 5e). The oxygen evolution results suggest *S. microadriaticum* cells are not stressed at pH 4 (Fig. 5f), and because there is still EET at pH 4, it suggests EET is an indicator of cell health. At pH 10, however, while steady state photocurrent levels were still positive (Fig. 5e), oxygen production was low (Fig. 5f). This further confirms uncoupling between EET and photosynthesis. However, it is worth noting that oxygen solubility can decrease with increasing ionic strength^49^, so addition of HCl and NaOH to adjust pH may have affected oxygen production measurements, especially at pH extremes.

To assess the predominance of the EET pathway under different stress conditions, we can calculate the proportion of electrons generated during photosynthesis that are directed to the EET pathway. The quantity of electrons generated by photosynthesis can be estimated from oxygen evolution measurements because for every mole of oxygen produced in water-splitting, 4 moles of electrons are generated. The number of moles of electrons that are funnelled through EET can be estimated from the charge of the photocurrent (see Methods section). Our hypothesis was that as cells experience redox stress, a higher percentage of electrons would be directed to EET to prevent the formation of ROS inside the cell. We found that a higher percentage was exported at alkaline pHs (Supplementary Fig. 6c). However, between 25°C and 40°C, the percentage remained relatively constant (Supplementary Fig. 6d). A higher percentage of electrons was exported in the regime of decreasing light compared to the regime of increasing light, but a lower overall percentage of electrons was exported at high light intensities (Supplementary Fig. 6e). We predict that alternative sinks would be activated in high light, so a smaller percentage of electrons would go through the EET pathway. Light stress and high temperature are both known to induce ROS production, and dinoflagellate algae are more suited to acidic or neutral pHs. Taken together, this suggests that *S. microadriaticum* can alter the flux of electrons into EET to adapt to stressful environmental conditions, but that the cells do not simply increase EET activity in stressful conditions to limit ROS production.

Overall, the results demonstrate that the electrochemical platform can give a readout of bioenergetics in stress conditions linked to bleaching. The evidence for uncoupling between EET and photosynthesis suggests that photocurrents are not simply a measure of photosynthetic activity, but rather are a reflection of the overall redox balance in the cell.

## Discussion

In this study, we developed an electrochemical platform to measure EET in *S. microadriaticum*. We confirm that EET activity can be measured, that it can be applied to understand intracellular bioenergetics, and that it is responsive to stressors involved in coral bleaching.

This observation of EET from dinoflagellate algae opens up fundamental questions relating to the interactions between dinoflagellate algae and corals during symbiosis. Interspecies EET has been reported before in microbial communities^50,51^, and electromechanical communication between bacteria and eukaryotes has been demonstrated in engineered living materials^52^. However, to the best of our knowledge EET has not been reported between symbionts and their animal hosts. Although we have not demonstrated directly here that dinoflagellates in their symbiotic state can perform EET, we have shown that they do in their free-living state. Assuming they do so in their symbiont state as well (which seems likely, since they remain photosynthetically active), EET is likely to interact with the coral host cell, given that the symbionts are contained in a confined space within coral cells (the symbiosome^53^, Fig. 1c), and that the electron carrier produced by *S. microadriaticum* is diffusible (Fig. 4b-c). The symbiosome is decorated with receptors^54^ that could transfer electron carriers across the membrane and into the coral cell. There is little data in the literature on the metabolites exchanged between dinoflagellates and corals (often called ‘photosynthate’), let alone ones with redox activity. Therefore, this technique could complement metabolomics studies^55–58^, which have thus far focused mainly on carbohydrates and lipids^59^.

If symbiotic dinoflagellate algae perform EET, then this pathway could have several functions in the symbiosis. First, EET could be used to share energy with the coral host by means of reducing equivalents that funnel out excess energy from photosynthesis (Fig. 5a). This way, EET would benefit both corals and dinoflagellates, for which it could also act as a photoprotection pathway. Alternatively, EET may be involved in the initiation of bleaching by acting as a stress signalling pathway between dinoflagellate algae and coral hosts. For example, an increase in EET activity could be used by dinoflagellates to signal redox stress to their host before ROS-mediated damage can occur. This could explain the higher steady state photocurrent measured in high light (Fig. 5a). High visible light can trigger bleaching by itself, and it can also accentuate the effect of heat stress by causing photoinhibition^3,60–62^. Lastly, EET activity could be used to quench ROS, thus preventing ROS from reacting with coral proteins. At high temperature, the loss of EET activity (Fig. 5c) and its associated antioxidant activity could then trigger bleaching. This is supported by a previous evidence of stress markers produced by *S. microadriaticum* at 32°C, including ROS^63^, nitric oxide^64^, dimethylsulfoniopropionate^65^ and heat shock proteins^66^.

The dependency of EET on intracellular bioenergetics and environmental conditions could make the electrochemical platform developed here an attractive system for coral diagnostics or assaying, if it can be adapted to dinoflagellate algae living inside of coral. In contrast to current coral biosensors which monitor photosynthetic activity with fluorimetry^67,68^, the electrochemical platform developed here could provide a readout of whole-cell redox balance (Fig 5). These measurements would be particularly valuable, since many pathways other than photosynthesis can be involved in bleaching^3^. EET measurements could be used as a proxy for cell health, similar to how the bioelectrochemical literature uses measurements of EET instead of growth rate because it provides better temporal resolution and reflects metabolic activity rather than optical density^11,69^. Finally, electrochemical techniques are easily scalable, and form the basis of many biosensors^70^. This makes the platform developed here an attractive system for low-cost coral diagnostics or other biotechnology developments.

Overall, we demonstrated that *S. microadriaticum* performs EET, a feature which could be common to the wider group of dinoflagellate algae. The electrochemical platform developed here will allow researchers to explore fundamental questions about the coral-dinoflagellate symbiosis, and work towards building diagnostic tools for corals to mitigate the bleaching crisis.

## Methods

### Biological strains and culturing

Artificial seawater was prepared in MilliQ® water by dissolving Coral Pro Salt (33.6 g L^-1^, Red Sea) and tricine (0.5 g L^-1^, ≥99%, 5704-04-1, Sigma-Aldrich), adjusting the pH to 7.8 with NaOH, and autoclaving. F/2 medium was made by adding a filter-sterilised f/2 supplement stock (0.15% v/v, Phytoplankton Nutrient Guillard’s F/2 Medium, Reef Phyto) to the artificial seawater medium. Wild-type *Symbiodinium microadriaticum* (strain CCMP2467) was cultured in liquid f/2 medium, without shaking, at 26°C in a diel cycle of 14 h light and 10 h dark. The light intensity used was 20 μmol photons m^-2^ s^-1^ of photosynthetically active radiation. Cell counts were performed with a haemocytometer (Z359629-1EA, Merck). UV-vis spectrophotometry was performed with a Cary 60 instrument (Agilent Technologies, USA). Pigments were extracted with 0.5 M ammonium acetate and methanol as described previously^71^.

### Three-electrode system fabrication

Working electrodes were made by cutting fluorine-doped tin oxide (FTO)-coated glass (735167-1EA, Sigma Aldrich) into 3 x 1.2 cm rectangles. To prepare the mesoITO substrate, the FTO-coated glass was cleaned by sonication at 37 kHz first in 99% isopropanol, then 99% ethanol. ITO nanoparticles (99.5%, 50926-11-9, ThermoFisher) were solubilised in acetic acid (5M) in ethanol to produce a 20% w/v solution which was sonicated for 30 min on ice. Circular templates of 1 cm^2^ geometric area were prepared from adhesive tape and stuck onto the FTO-coated glass. The solubilised ITO was spread over the circular templates and scraped with a glass coverslip to obtain a thin layer of ITO (∼ 2.5 μm). Once dried, the templates were removed and the electrodes were dried in a furnace (Carbolite ELF 11/14B/301) at 450°C for 20 min, at a ramp rate of 4°C min^-1^. Platinum mesh counter electrodes were made by attaching 1 cm x 0.5 cm segments of platinum mesh (Goodfellow, UK) to tin-coated copper wires of diameter 1 mm. Ag/AgCl reference electrodes were made in-house and store in a saturated KCl solution. The reference electrodes were calibrated with a commercial Ag/AgCl electrode (MF-2052, BASi). To build the three-electrode setup, a custom-made polyether ether ketone dish with an opening at the bottom was fabricated. The dish was filled with 5 mL f/2 electrolyte, and *S. microadriaticum* cells were added directly onto the working electrode using a micropipette.

### Chronoamperometry

Chronoamperometry was performed using a CompactStat.h potentiostat (Ivium Technologies). A three-electrode system composed of a working mesoITO electrode, an Ag/AgCl reference and a platinum mesh counter were used. Unless specified otherwise, measurements were performed at room temperature; with chopped 680 nm light (60 s light, 90 s dark) at 50 μmol photons m^-2^ s^-1^; 3 x 10^6^ cells; and a potential of 0.3 V vs SHE. A pre-equilibration time of 90 s was used before each measurement. For stepped chronoamperometry, the potential was varied from 0.1 V to 0.6 V vs SHE. Where applicable, 3-(3,4-Dichloro-phenl)-1,1-dimethylurea (DCMU, ≥98%, D2425, Merck) or rotenone (≥95%, 83-79-4, Sigma Aldrich) were solubilised in DMSO and added to the electrolyte. SHAM (99%, 11424533, Thermo Scientific) was solubilised in f/2.

### Scanning electrochemical microscopy

Scanning electrochemical microscopy was performed using a CH920C scanning electrochemical microscope (CH Instruments, USA). A three-electrode system composing a platinum wire (CH Instruments, USA), Ag/AgCl reference electrode (ALS, Japan) and a 10 µm diameter platinum UME with an RG value of 6 (CH Instruments, USA) were used. The UME probe was polished with 0.05 µm 𝛾-alumina prior to use. After 2 h of incubation in the light or in the dark, the following measurements were taken: a CV of the bulk 300 µm above the surface (50 mV s^-1^), a probe approach curve at 1 µm s-^1^ and 0.7 V vs SHE with 5 min pre-equilibration time, and CVs 20 µm above the surface (50 mV s^-1^). Control CVs were performed at 10 mV s^-1^ with a sample interval of 1 mV. All experiments were performed at room temperature with, where applicable, 100 μmol photons m^-2^ s^-1^ of 680 nm light.

### Stress experiments

For light stress experiments, *S. microadriaticum* cells were kept in the dark for 1 h and subsequently exposed to a series of light intensities, ranging from 50 to 1000 μmol photons m^-2^ s^-1^. In the light stress regime, the light intensities were used in decreasing order. For temperature stress experiments, temperature was increased at a ramp rate of 0.5°C min^-1^ using a hot plate (for electrochemistry) or water bath (for oxygen evolution). At each measured temperature, the conditions were held for 15 min while chronoamperometry was performed. For pH stress experiments, f/2 was adjusted to the desired pH using HCl or NaOH. *S. microadriaticum* cells were centrifuged at 10,000 g for 3 min and resuspended directly into the measured pH. All oxygen evolution rates were measured using an oxygen sensor (OXROB10-HS, Pyroscience) connected to a multi-analyte meter (FSPRO-2, Pyroscience).

### Data analysis

Steady state current intensities I_light_ and I_dark_ were calculated by averaging the last 5s of the current in the light and dark respectively. Steady state photocurrent was calculated by subtracting I_dark_ from I_light_. Potentials measured vs Ag/AgCl were converted to potentials vs SHE with the following conversion method: E_SHE_ = E_Ag/AgCl_ + 0.209 V.

The number of moles of electrons detected electrochemically was calculated by:

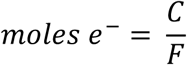

where *C* is charge in A · s and *F* is the Faraday constant in C mol^-1^.

The number of moles of electrons produced through photosynthesis was calculated by:

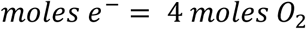

The percentage of electrons directed to the EET pathway from photosynthesis was calculated by:

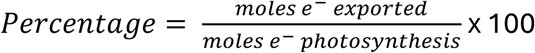

All the data analysis was performed using in-house Python scripts, which can be found on GitHub (https://github.com/Zhang-Lab-Cambridge/DinoBot.git).

## Acknowledgments

The authors are grateful for support from the Natural Environmental Research Council (2894423 to L.M. and NE/X010503/1 to A.C.B., R.E.R.N., and C.J.H.) and Trinity Hall, Cambridge (to J.M.L.). S.M. gratefully acknowledges funding by the Royal Society University Research Fellowship (URF, URF\R1\231513) and the support funding the SECM purchase from the Henry Royce Institute for Advanced Materials, Cambridge. We thank Dr Linying Shang, Robin Scullion, Leanne Milburn and Dr Leonid Digel for their help in reviewing the manuscript. We thank Prof. Patrick Unwin for his invaluable suggestions for the SECM controls.

## Author contributions

All authors contributed to designing experiments. J.T.S. and L.M. performed SECM experiments, S.M. provided the SECM expertise and equipment. L.M. performed all other experiments. A.C.B. assisted with cell culturing. J.M.L., J.T.S., A.C.B., R.E.R.N., and C.J.H. assisted in data interpretation and manuscript editing. L.M. and J.Z.Z. wrote the manuscript. R.E.R.N., C.J.H. and J.Z.Z. conceptualised the project initially.

## References

1 Muscatine, L., Porter, J. W. & Kaplan, I. R. Resource partitioning by reef corals as determined from stable isotope composition. Mar. Biol. 100, 185–193 (1989). 10.1007/bf00391957

2 Davy, S. K., Allemand, D. & Weis, V. M. Cell biology of cnidarian-dinoflagellate symbiosis. Microbiol. Mol. Biol. Rev. 76, 229–261 (2012). 10.1128/mmbr.05014-11

3 Helgoe, J., Davy, S. K., Weis, V. M. & Rodriguez-Lanetty, M. Triggers, cascades, and endpoints: connecting the dots of coral bleaching mechanisms. Biol. Rev. Camb. Philos. Soc. 9, 1–38 (2024). 10.1111/brv.13042

4 Mellin, C. et al. Cumulative risk of future bleaching for the world’s coral reefs. Sci. Adv. 10, eadn9660 (2024). 10.1126/sciadv.adn9660

5 Chen, J. E., Barbrook, A. C., Cui, G., Howe, C. J. & Aranda, M. The genetic intractability of Symbiodinium microadriaticum to standard algal transformation methods. PLoS One 14, e0211936 (2019). 10.1371/journal.pone.0211936

6 Gornik, S. G. et al. Nuclear transformation of a dinoflagellate symbiont of corals. Frontiers in Marine Science 9, 1035413 (2022). 10.3389/fmars.2022.1035413

7 Nimmo, I. C. et al. Genetic transformation of the dinoflagellate chloroplast. Elife 8, e45292 (2019). 10.7554/eLife.45292

8 Gornik, S. G. et al. Loss of nucleosomal DNA condensation coincides with appearance of a novel nuclear protein in dinoflagellates. Curr. Biol. 22, 2303–2312 (2012). 10.1016/j.cub.2012.10.036

9 Howe, C. J. & Barbrook, A. C. Dinoflagellate chloroplasts as a model for extreme genome reduction and fragmentation in organelles - The COCOA principle for gene retention. Protist 175, 126048 (2024). 10.1016/j.protis.2024.126048

10 Lovley, D. R. & Holmes, D. E. Electromicrobiology: the ecophysiology of phylogenetically diverse electroactive microorganisms. Nat. Rev. Microbiol. 20, 5–19 (2022). 10.1038/s41579-021-00597-6

11 Digel, L., Bonné, R. & Aiyer, K. Are all microbes electroactive? Cell Rep. Phys. Sci. 5, 102200 (2024). 10.1016/j.xcrp.2024.102200

12 Wey, L. T. et al. The development of biophotovoltaic systems for power generation and biological analysis. ChemElectroChem 6, 5375–5386 (2019). 10.1002/celc.201900997

13 Gupta, D., Chen, K., Elliott, S. J. & Nayak, D. D. MmcA is an electron conduit that facilitates both intracellular and extracellular electron transport in Methanosarcina acetivorans. Nat. Commun. 15, 3300 (2024). 10.1038/s41467-024-47564-2

14 Lovley, D. R. Syntrophy Goes Electric: Direct Interspecies Electron Transfer. Annu. Rev. Microbiol. 71, 643–664 (2017). 10.1146/annurev-micro-030117-020420

15 Saper, G. et al. Live cyanobacteria produce photocurrent and hydrogen using both the respiratory and photosynthetic systems. Nat. Commun. 9, 2168 (2018). 10.1038/s41467-018-04613-x

16 Kusama, S. et al. Order-of-magnitude enhancement in photocurrent generation of Synechocystis sp. PCC 6803 by outer membrane deprivation. Nat. Commun. **13**, 3067 (2022). 10.1038/s41467-022-30764-z

17 Bombelli, P. et al. Quantitative analysis of the factors limiting solar power transduction by Synechocystis sp. PCC 6803 in biological photovoltaic devices. Energ. Environ. Sci. **4**, 4690-4698 (2011). 10.1039/C1EE02531G

18. Szabó, M., Larkum, A. W. D. & Vass, I. in Photosynthesis in Algae: Biochemical and Physiological Mechanisms (eds Anthony W. D. Larkum, Arthur R. Grossman, & John A. Raven) 459-488 (Springer International Publishing, 2020).

19 Lesser, M. P. Elevated temperatures and ultraviolet radiation cause oxidative stress and inhibit photosynthesis in symbiotic dinoflagellates. Limnol. Oceanogr. 41, 271–283 (1996). 10.4319/lo.1996.41.2.0271

20 Warner, M. E., Fitt, W. K. & Schmidt, G. W. Damage to photosystem II in symbiotic dinoflagellates: A determinant of coral bleaching. Proc. Natl. Acad. Sci. USA 96, 8007–8012 (1999). 10.1073/pnas.96.14.8007

21 Rosic, N., Delamare-Deboutteville, J. & Dove, S. Heat stress in symbiotic dinoflagellates: Implications on oxidative stress and cellular changes. Sci. Total Environ. 944, 173916 (2024). 10.1016/j.scitotenv.2024.173916

22 Rehman, A. U. et al. Symbiodinium sp. cells produce light-induced intra- and extracellular singlet oxygen, which mediates photodamage of the photosynthetic apparatus and has the potential to interact with the animal host in coral symbiosis. New Phytol. 212, 472–484 (2016). 10.1111/nph.14056

23 Slavov, C. et al. "Super-quenching" state protects Symbiodinium from thermal stress - Implications for coral bleaching. Biochim. Biophys. Acta 1857, 840–847 (2016). 10.1016/j.bbabio.2016.02.002

24 LaJeunesse, T. C. Validation and description of Symbiodinium microadriaticum, the type species of Symbiodinium (Dinophyta). J. Phycol. 53, 1109–1114 (2017). 10.1111/jpy.12570

25 Kato, M., Cardona, T., Rutherford, A. W. & Reisner, E. Photoelectrochemical water oxidation with photosystem II integrated in a mesoporous indium–tin oxide electrode. J. Am. Chem. Soc. 134, 8332–8335 (2012). 10.1021/ja301488d

26 Niedzwiedzki, D. M., Jiang, J., Lo, C. S. & Blankenship, R. E. Spectroscopic properties of the Chlorophyll a–Chlorophyll c 2–Peridinin-Protein-Complex (acpPC) from the coral symbiotic dinoflagellate Symbiodinium. Photosynthesis Research 120, 125–139 (2014). 10.1007/s11120-013-9794-5

27 Brewer, S. H. & Franzen, S. Calculation of the electronic and optical properties of indium tin oxide by density functional theory. Chem. Phys. 300, 285–293 (2004). 10.1016/j.chemphys.2003.11.039

28 Loeblich, A. R. & Sherley, J. L. Observations on the theca of the motile phase of free-living and symbiotic isolates of Zooxanthella microadriatica (Freudenthal) comb.nov. J. Mar. Biolog. Assoc. U.K. 59, 195–205 (1979). 10.1017/S0025315400046270

29 Tortorelli, G. et al. Cell surface carbohydrates of symbiotic dinoflagellates and their role in the establishment of cnidarian–dinoflagellate symbiosis. ISME J. 16, 190–199 (2022). 10.1038/s41396-021-01059-w

30 Vicente-Garcia, C. et al. Living diatom microalgae for desiccation-resistant electrodes in biophotovoltaic devices. ACS Sustainable Chemistry & Engineering 12, 11120–11129 (2024). 10.1021/acssuschemeng.4c00935

31 Zhang, J. Z. et al. Competing charge transfer pathways at the photosystem II– electrode interface. Nat. Chem. Biol. 12, 1046–1052 (2016). 10.1038/nchembio.2192

32 Aihara, Y., Takahashi, S. & Minagawa, J. Heat induction of cyclic electron flow around photosystem I in the symbiotic dinoflagellate Symbiodinium. Plant Physiol. 171, 522–529 (2016). 10.1104/pp.15.01886

33 Fransolet, D., Roberty, S. & Plumier, J.-C. Impairment of symbiont photosynthesis increases host cell proliferation in the epidermis of the sea anemone Aiptasia pallida. Mar. Biol. 161, 1735–1743 (2014). 10.1007/s00227-014-2455-1

34 Zhao, L.-S. et al. Architecture of symbiotic dinoflagellate photosystem I–light-harvesting supercomplex in Symbiodinium. Nat. Commun. 15, 2392 (2024). 10.1038/s41467-024-46791-x

35 Oakley, C., Hopkinson, B. & Schmidt, G. Mitochondrial terminal alternative oxidase and its enhancement by thermal stress in the coral symbiont Symbiodinium. Coral Reefs 33, 543–522 (2014). 10.1007/s00338-014-1147-0

36 Danne, J. C., Gornik, S. G., MacRae, J. I., McConville, M. J. & Waller, R. F. Alveolate Mitochondrial Metabolic Evolution: Dinoflagellates Force Reassessment of the Role of Parasitism as a Driver of Change in Apicomplexans. Molecular Biology and Evolution 30, 123–139 (2012). 10.1093/molbev/mss205

37 Butterfield, E. R., Howe, C. J. & Nisbet, R. E. R. An analysis of dinoflagellate metabolism using EST data. Protist 164, 218–236 (2013). 10.1016/j.protis.2012.09.001

38 Raven, J. A. & Beardall, J. Consequences of the genotypic loss of mitochondrial Complex I in dinoflagellates and of phenotypic regulation of Complex I content in other photosynthetic organisms. J. Exp. Bot. 68, 2683–2692 (2017). 10.1093/jxb/erx149

39 Pierangelini, M., Thiry, M. & Cardol, P. Different levels of energetic coupling between photosynthesis and respiration do not determine the occurrence of adaptive responses of Symbiodiniaceae to global warming. New Phytol. 228, 855–868 (2020). 10.1111/nph.16738

40 McKelvey, K., Martin, S., Robinson, C. & Unwin, P. R. Quantitative local photosynthetic flux measurements at isolated chloroplasts and thylakoid membranes using scanning electrochemical microscopy (SECM). J. Phys. Chem. B 117, 7878–7888 (2013). 10.1021/jp403048f

41 Chen, B., Hu, Q., Xiong, Q., Zhang, F. & He, P. An ultrasensitive scanning electrochemical microscopy (SECM)-based DNA biosensing platform amplified with the long self-assembled DNA concatemers. Electrochim. Acta 192, 127–132 (2016). 10.1016/j.electacta.2015.12.102

42 Su, L., Tong, Y., Shu, T., Gong, W. & Zhang, X. Single-walled carbon nanotube ensembles modified gold ultramicroelectrodes prepared by self-assembly deposition method with 1-(1-pyrenyl)-1-methanethiol monolayer as an adhesion layer. Electrochem. Commun. 20, 163–166 (2012). 10.1016/j.elecom.2012.04.019

43 Arbenin, A. Y. et al. Prospects of application of ultramicroelectrode ensembles for voltammetric determination of compounds with close standard electrode potentials and different diffusion coefficients. Chemosensors 10, 433 (2022). 10.3390/chemosensors10100433.

44 Wu, Y., Ma, P., Liu, Y. & Li, S. Diffusion coefficients of l-proline, l-threonine and l-arginine in aqueous solutions at 25°C. Fluid Ph. Equilib. 186, 27–38 (2001). 10.1016/S0378-3812(01)00355-7

45 Jia, S. et al. Comparison of physiological and transcriptome responses of corals to strong light and high temperature. Ecotoxicol. Environ. Saf. 273, 116143 (2024). 10.1016/j.ecoenv.2024.116143

46 Hughes, T. P. et al. Spatial and temporal patterns of mass bleaching of corals in the Anthropocene. Science 359, 80–83 (2018). 10.1126/science.aan8048

47 Langdon, C. & Atkinson, M. J. Effect of elevated pCO2 on photosynthesis and calcification of corals and interactions with seasonal change in temperature/irradiance and nutrient enrichment. J. Geophys. Res. Oceans 110, C09S07 (2005). 10.1029/2004JC002576

48 Anthony, K. R., Kline, D. I., Diaz-Pulido, G., Dove, S. & Hoegh-Guldberg, O. Ocean acidification causes bleaching and productivity loss in coral reef builders. Proc. Natl. Acad. Sci. USA 105, 17442–17446 (2008). 10.1073/pnas.0804478105

49. Xing, W. et al. in Rotating Electrode Methods and Oxygen Reduction Electrocatalysts (eds Wei Xing, Geping Yin, & Jiujun Zhang) 1-31 (Elsevier, 2014).

50 Wang, W. & Sheng, Y. Interactions between Microcystis and its associated bacterial community on electron transfer and transcriptomic processes. Sci. Total Environ. 950, 175372 (2024). 10.1016/j.scitotenv.2024.175372

51 Nagarajan, H. et al. Characterization and modelling of interspecies electron transfer mechanisms and microbial community dynamics of a syntrophic association. Nat. Commun. 4, 2809 (2013). 10.1038/ncomms3809

52 Atkinson, J. T. Inter-kingdom electromechanical communication. Nat. Chem. Biol. 20, 1250–1251 (2024). 10.1038/s41589-024-01687-1

53 Wakefield, T. S. & Kempf, S. C. Development of host- and symbiont-specific monoclonal antibodies and confirmation of the origin of the symbiosome membrane in a cnidarian–dinoflagellate symbiosis. Biol. Bull. 200, 127–143 (2001). 10.2307/1543306

54 Thies, A. B., Quijada-Rodriguez, A. R., Zhouyao, H., Weihrauch, D. & Tresguerres, M. A Rhesus channel in the coral symbiosome membrane suggests a novel mechanism to regulate NH(3) and CO(2) delivery to algal symbionts. Sci. Adv. 8, eabm0303 (2022). 10.1126/sciadv.abm0303

55 Hillyer, K. E., Tumanov, S., Villas-Bôas, S. & Davy, S. K. Metabolite profiling of symbiont and host during thermal stress and bleaching in a model cnidarian– dinoflagellate symbiosis. J. Exp. Biol. 219, 516–527 (2016). 10.1242/jeb.128660

56 Roach, T. N. F. et al. Metabolomic signatures of coral bleaching history. *Nat*. Ecol. Evol. 5, 495–503 (2021). 10.1038/s41559-020-01388-7

57 Williams, A. et al. Metabolomic shifts associated with heat stress in coral holobionts. Sci. Adv. 7, eabd4210 (2021). 10.1126/sciadv.abd4210

58 Matthews, J. L., Bartels, N., Elahee Doomun, S. N., Davy, S. K. & De Souza, D. P. Gas chromatography-mass spectrometry-based targeted metabolomics of hard coral samples. J. Vis. Exp., e65628 (2023). 10.3791/65628

59 Rosset, S. L. et al. The molecular language of the cnidarian–dinoflagellate symbiosis. Trends Microbiol. 29, 320–333 (2021). 10.1016/j.tim.2020.08.005

60 Takahashi, S. & Murata, N. How do environmental stresses accelerate photoinhibition? Trends Plant Sci. 13, 178–182 (2008). 10.1016/j.tplants.2008.01.005

61 Tagliafico, A., Baker, P., Kelaher, B., Ellis, S. & Harrison, D. The effects of shade and light on corals in the context of coral bleaching and shading technologies. Frontiers in Marine Science 9, 919382 (2022). 10.3389/fmars.2022.919382

62 Berg, J. T., David, C. M., Gabriel, M. M. & Bentlage, B. Fluorescence signatures of persistent photosystem damage in the staghorn coral Acropora cf. pulchra (Anthozoa: Scleractinia) during bleaching and recovery. *Mar*. Biol. Res. 16, 643–655 (2020). 10.1080/17451000.2021.1875245

63 Tchernov, D. et al. Membrane lipids of symbiotic algae are diagnostic of sensitivity to thermal bleaching in corals. Proc. Natl. Acad. Sci. USA 101, 13531–13535 (2004). 10.1073/pnas.0402907101

64 Bouchard, J. N. & Yamasaki, H. Heat stress stimulates nitric oxide production in Symbiodinium microadriaticum: a possible linkage between nitric oxide and the coral bleaching phenomenon. Plant Cell Physiol. 49, 641–652 (2008). 10.1093/pcp/pcn037

65 McLenon, A. L. & DiTullio, G. R. Effects of increased temperature on dimethylsulfoniopropionate (DMSP) concentration and methionine synthase activity in Symbiodinium microadriaticum. Biogeochemistry 110, 17–29 (2012). 10.1007/s10533-012-9733-0

66 Oakley, C. A., Newson, G. I., Peng, L. & Davy, S. K. The symbiodinium proteome response to thermal and nutrient stresses. Plant Cell Physiol. 64, 433–447 (2023). 10.1093/pcp/pcac175

67 Wangpraseurt, D., Lichtenberg, M., Jacques, S. L., Larkum, A. W. D. & Kühl, M. Optical properties of corals distort variable chlorophyll fluorescence measurements. Plant Physiol. 179, 1608–1619 (2019). 10.1104/pp.18.01275

68 Beer, S., Ilan, M., Eshel, A., Weil, A. & Brickner, I. Use of pulse amplitude modulated (PAM) fluorometry for in situ measurements of photosynthesis in two Red Sea faviid corals. Mar. Biol. 131, 607–612 (1998). 10.1007/s002270050352

69 Grobbler, C. et al. Effect of the anode potential on the physiology and proteome of Shewanella oneidensis MR-1. Bioelectrochemistry 119, 172–179 (2018). 10.1016/j.bioelechem.2017.10.001

70 Wu, J., Liu, H., Chen, W., Ma, B. & Ju, H. Device integration of electrochemical biosensors. Nat. Rev. Bioeng. 1, 346–360 (2023). 10.1038/s44222-023-00032-w

71 Rogers, J. E. & Marcovich, D. A simple method for the extraction and quantification of photopigments from Symbiodinium spp. J. Exp. Mar. Biol. Ecol. 353, 191–197 (2007). 10.1016/j.jembe.2007.08.022

