## Supplementary figures for "Bioelectricity generation by *Symbiodinium microadriaticum*: a symbiont-forming photosynthetic dinoflagellate alga from coral reefs"

### Supplementary information

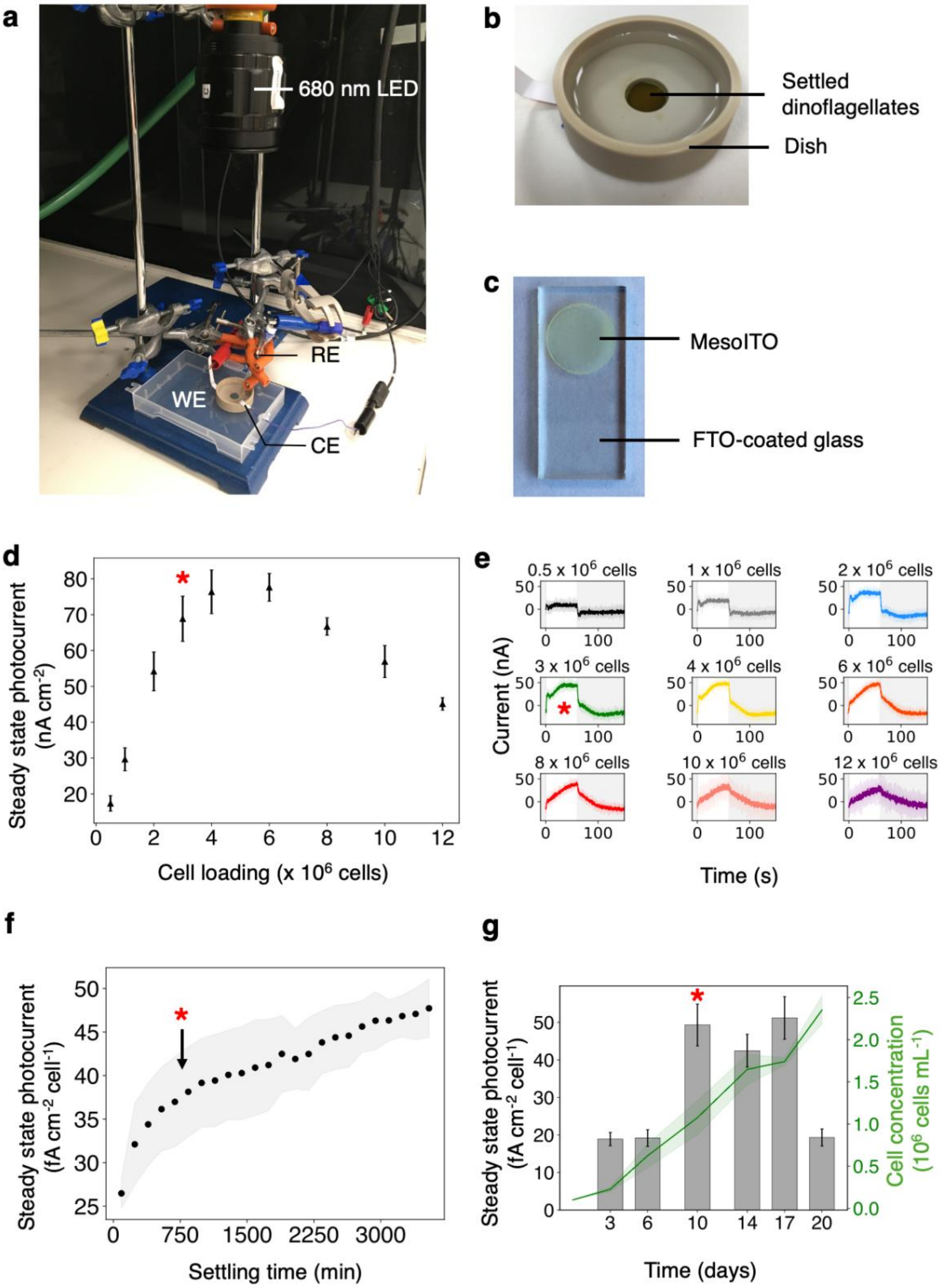
a
680 nm LED
RE
WE
CE
b
Settled dinoflagellates
Dish
c
MesoITO
FTO-coated glass
d
Steady state photocurrent (nA cm-2)
Cell loading (x 10^6 cells)
e
0.5 x 10^6 cells
1 x 10^6 cells
2 x 10^6 cells
3 x 10^6 cells
4 x 10^6 cells
6 x 10^6 cells
8 x 10^6 cells
10 x 10^6 cells
12 x 10^6 cells
Current (nA)
Time (s)
f
Steady state photocurrent (fA cm-2 cell-1)
Settling time (min)
g
Steady state photocurrent (fA cm-2 cell-1)
Cell concentration (10^6 cells mL-1)
Time (days)

**Supplementary Fig. 1: Optimisation of the electrochemical platform. a** Image of the electrochemical platform. **b** Electrochemical dish with *S. microadriaticum* cells settled onto the working electrode. **c** Working electrode comprised of a mesoITO substrate on FTO-coated glass. **d** Effect of cell loading on steady state photocurrent. **e** Effect of cell loading on photocurrent profile. **f** Steady state photocurrent over time as *S. microadriaticum* cells settle onto the working electrode. $T_0$ corresponds to the time the cells were added into the electrochemical dish. **g** Effect of growth stage on steady state photocurrent. Data shown are averages of three biological replicates containing five technical replicates each, error bars and shaded areas represent the standard error of the mean. Unless specified, measurements were obtained at 0.3 V vs SHE; with 680 nm light at 50 μmol photons $m^{-2}$ $s^{-1}$; and 3 x $10^6$ cells. Symbols (*) represent optimised conditions.

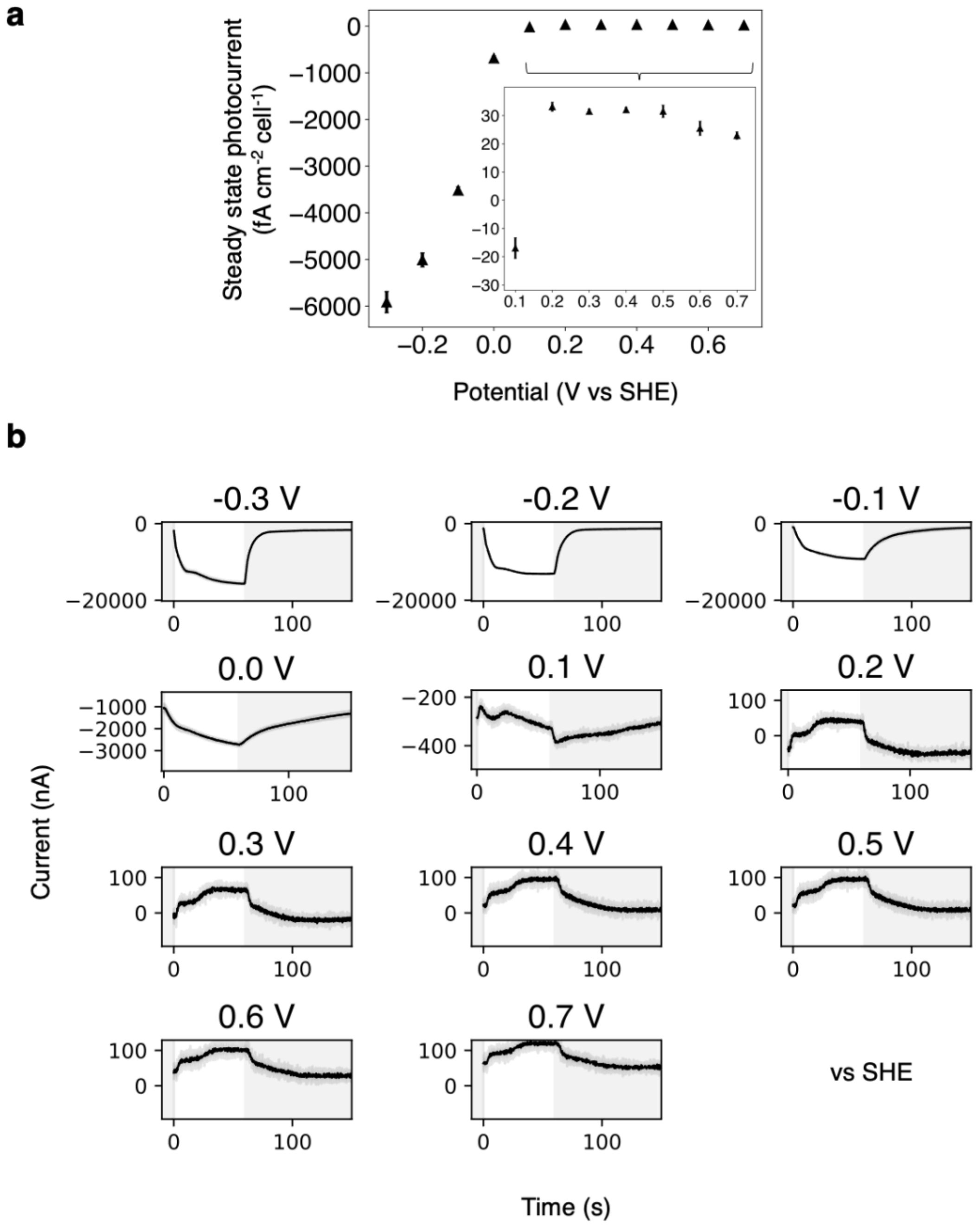

**Supplementary Fig. 2: Stepped chronoamperometry of S. microadriaticum cells on mesoITO. a** Steady state photocurrents obtained during stepped chronoamperometry from –0.3 V to 0.7 V vs SHE. Inset: steady state photocurrents in the 0.1 V to 0.7 V range. **b** Photocurrent profiles obtained from the stepped chronoamperometry. Data shown are averages of three biological replicates containing five technical replicates each, error bars represent the standard error of the mean and shaded areas represent standard deviation. Measurements were obtained with 680 nm light at 50 µmol photons $m^{-2}$ $s^{-1}$ and 3 x $10^6$ cells.

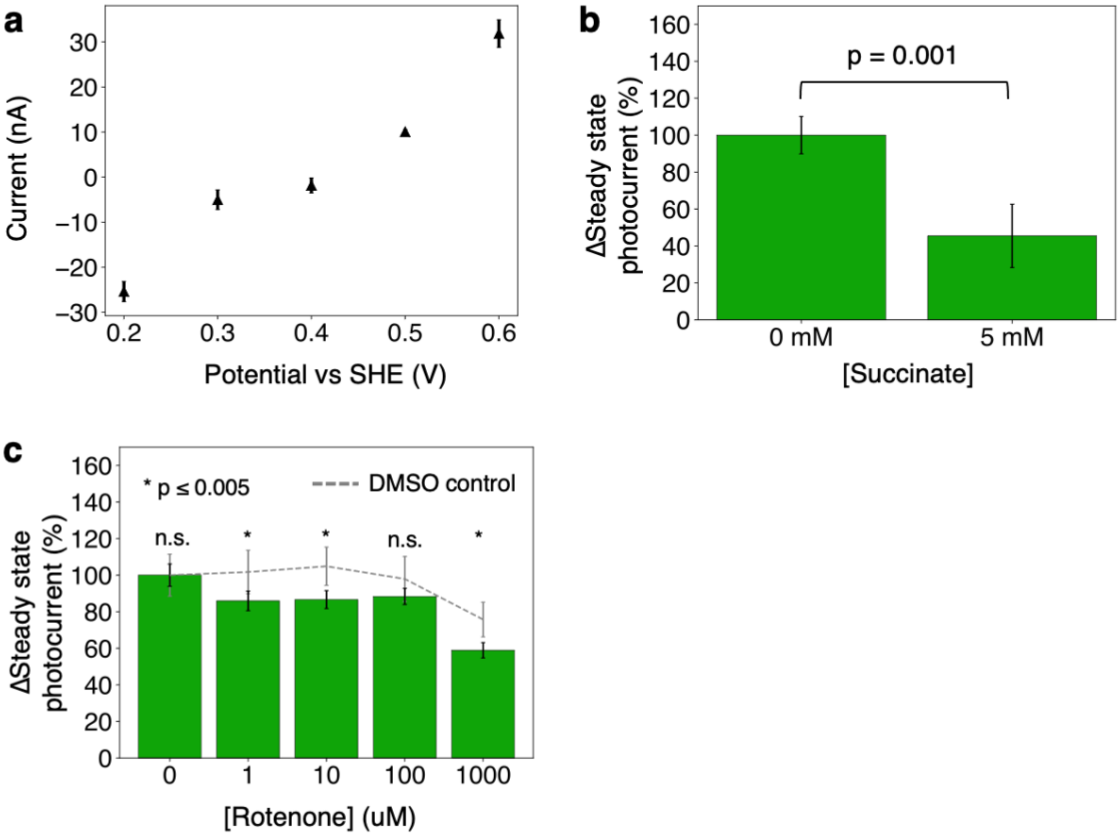

**Supplementary Fig. 3: Effect of respiratory activity on EET in S. microadriaticum. a** Dark currents obtained during stepped chronoamperometry from 0.2 V to 0.6 V vs SHE in f/2 medium without cells. **b** Effect of succinate on the steady state photocurrent. **c** Effect of rotenone on the steady state photocurrent. Data shown in *a* are averages of three biological replicates containing three technical replicates each. Data shown in *b* and *c* are averages of three biological replicates containing five technical replicates each. Error bars represent the standard error of the mean. Statistics were performed using a paired samples t-test. Unless specified, measurements were obtained at 0.3 V vs SHE; with 680 nm light at 50 µmol photons $m^{-2}$ $s^{-1}$; and 3 x $10^6$ cells.

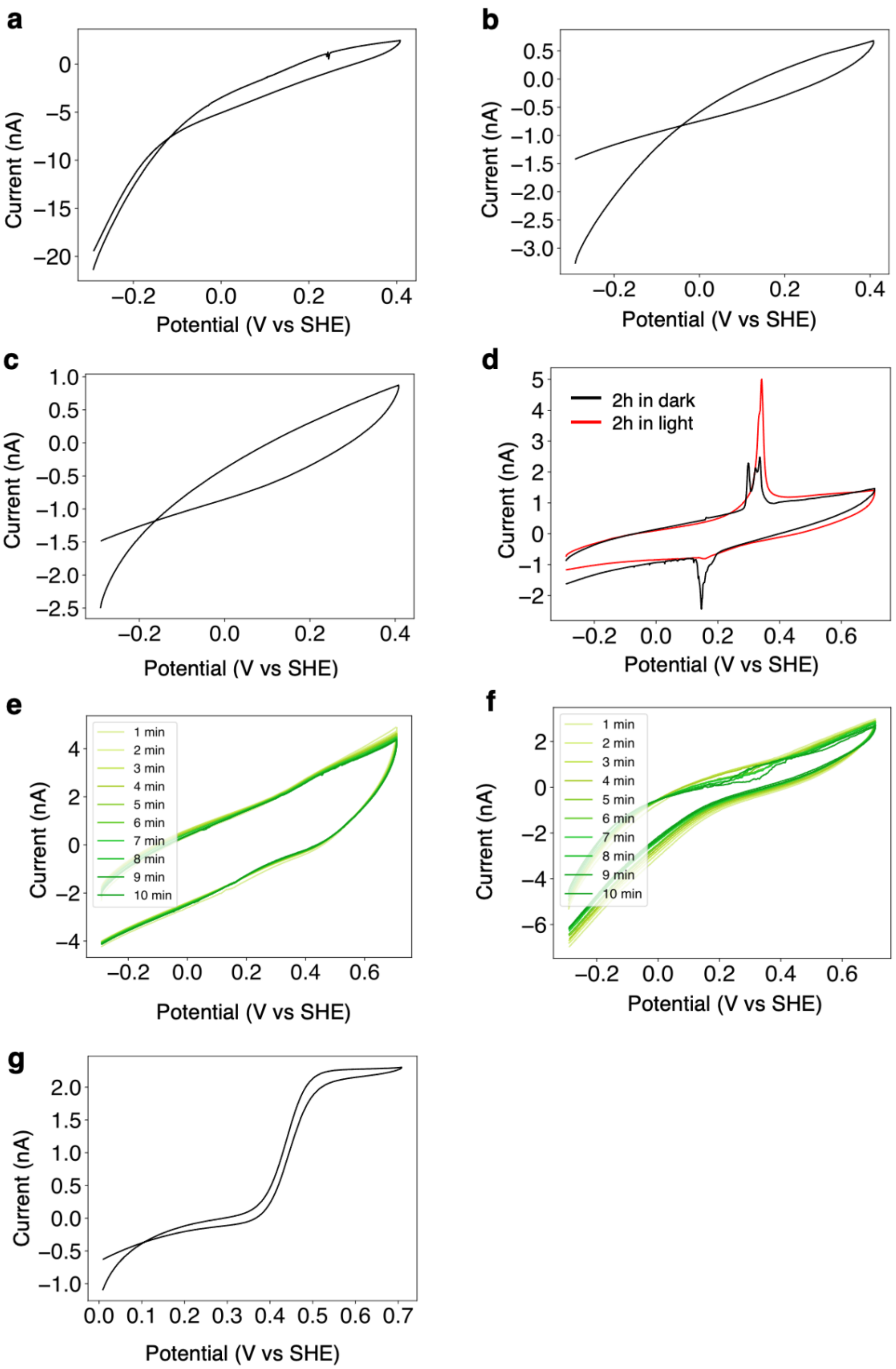
a
b
c
d
e
f
g
Current (nA)
Potential (V vs SHE)
2h in dark
2h in light
1 min
2 min
3 min
4 min
5 min
6 min
7 min
8 min
9 min
10 min

**Supplementary Fig. 4: Controls for SECM experiments.** **a** CV of f/2 medium. **b** Bulk CV of fresh f/2 medium to which *S. microadriaticum* cells were added. **c** CV of fresh f/2 medium to which *S. microadriaticum* cells were added, at the cell surface. **d** Average CVs of the bulk after 2h of incubation, in light or dark (n = 3). **e** Consecutive CVs of the surface with cells after 2 h of illumination and 100 μM DCMU. **f** Consecutive CVs of the surface without cells after 2 h of illumination. **g** CV of 1 mM FcMeOH mediator in 0.5 M NaCl solution. *a* - *c* and *g* were performed at 10 mV $s^{-1}$, the other CVs were performed at 50 mV $s^{-1}$ and a sample interval of 1 mV. Electrolyte was not purged with $N_2$.

At 25°C, the Randles-Sevcik equation is given as:

$$i_p = 2.69 \cdot 10^5 \cdot n^{\frac{3}{2}} AC\sqrt{D\nu}$$

With:

$i_p$ = maximum current in A

n = number of electrons involved in the redox process

A = electrode area in $cm^2$

C = concentration in mol $cm^{-3}$

D = diffusion coefficient in $cm^2$ $s^{-1}$

$\nu$ = scan rate in V $s^{-1}$

This can be rearranged to give:

$$C = \frac{i_p}{2.69 \cdot 10^5 \cdot n^{\frac{3}{2}} A\sqrt{D\nu}}$$

The following parameters were used:

Electrode surface area (A): 0.785 $cm^2$

Scan rate ($\nu$): 5 x $10^{-2}$ V $s^{-1}$

Diffusion coefficient ($D_0$): 5 x $10^6$ $cm^2$ $s^{-1}$

Maximum current ($i_p$): 25 x $10^{-9}$ A

Assuming a one-electron transfer process (n = 1), the estimated concentration of the diffusible electron carrier was:

$$C = \frac{i_p}{2.69 \cdot 10^5 \cdot n^{\frac{3}{2}} A\sqrt{D\nu}} = \frac{25 \cdot 10^{-9}}{2.69 \cdot 10^5 \cdot 1^{\frac{3}{2}} \cdot 0.785\sqrt{5 \cdot 10^6 \cdot 0.05}} = 0.237\, pM$$

Assuming a two-electron transfer process (n = 2), the estimated concentration of the diffusible electron carrier was:

$$C = \frac{i_p}{2.69 \cdot 10^5 \ \cdot n^{\frac{3}{2}} A\sqrt{D\nu}} = \frac{25 \cdot 10^{-9}}{2.69 \cdot 10^5 \ \cdot 2^{\frac{3}{2}} \ \cdot 0.785 \sqrt{5 \cdot 10^6 \ \cdot 0.05}} = 0.084\, pM$$

**Supplementary Fig. 5: Estimation of the concentration of the diffusible electron carrier using the Randles-Sevcik equation.**

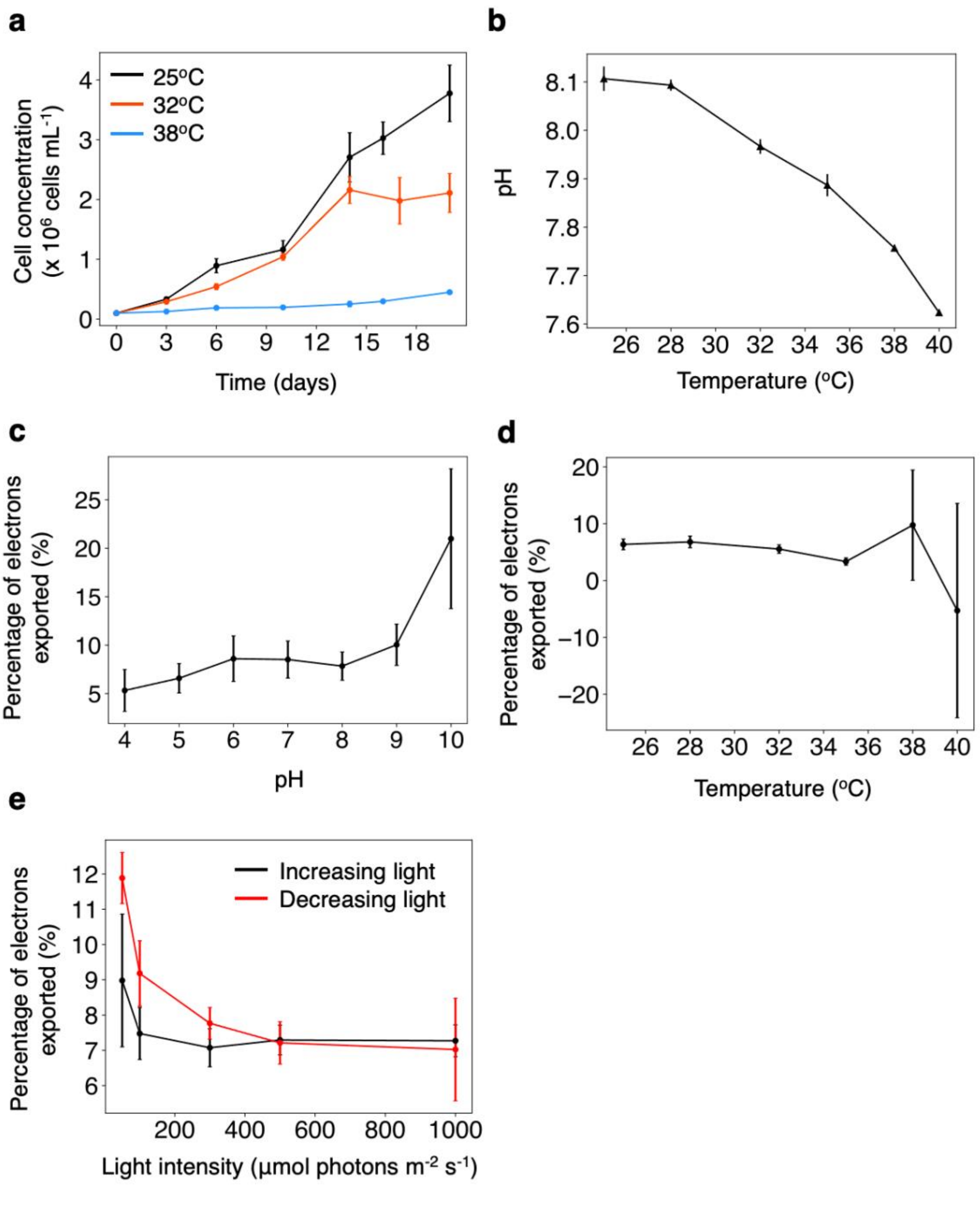

**Supplementary Fig. 6: Response of *S. microadriaticum* cells to different stress conditions. a** Effect of temperature on the growth of *S. microadriaticum*. **b** Effect of temperature on the pH of f/2 medium. **c-e** Percentage of electrons generated during photosynthesis that are directed to the EET pathway, calculated at different pHs (**c**), temperatures (**d**) and light intensities (**e**). See methods section for calculation details. Data shown are averages of three technical replicates, error bars represent standard deviation, except for *a* where they represent standard error of the mean.
